# Genetic risks of schizophrenia identified in a matched case-control study

**DOI:** 10.1101/702100

**Authors:** Kengo Oishi, Tomihisa Niitsu, Nobuhisa Kanahara, Yasunori Sato, Yoshimio Iwayama, Tomoko Toyota, Tasuku Hashimoto, Tsuyoshi Sasaki, Masayuki Takase, Takeo Yoshikawa, Masaomi Iyo

**Affiliations:** Department of Psychiatry, Chiba University Graduate School of Medicine, 1-8-1 Inohana, Chuou-ku, Chiba, Chiba 260-8670, Japan; Department of Preventive Medicine and Public Health, Keio University School of Medicine, 35 Shinanomachi Shinjuku-ku, Tokyo, 160-0016, Japan; Division of Medical Treatment and Rehabilitation, Chiba University Center for Forensic Mental Health, 1-8-1 Inohana, Chuou-ku, Chiba, Chiba 260-8670, Japan; Laboratory for Molecular Psychiatry, RIKEN Center for Brain Science, Wako, 351-0198 Saitama, Japan; Department of Child Psychiatry, Chiba University Hospital, 1-8-1 Inohana, Chuou-ku, Chiba, Chiba 260-8670, Japan

## Abstract

Genetic association studies of schizophrenia may be confounded by the pathological heterogeneity and multifactorial nature of this disease. We demonstrated previously that combinations of the three functional single nucleotide polymorphisms (SNPs) rs10770141 of tyrosine hydroxylase (TH) gene, rs4680 of catechol-O-methyltransferase (COMT) gene, and rs1800497 of dopamine D2 receptor (DRD2) gene may be associated with schizophrenia onset, and we tested those associations herein. Methods: We conducted a secondary study of 2,542 individuals in age- and sex-matched case-control populations. The schizophrenia diagnosis was based on the DSM-IV. To reduce the influence of confounders (age and sex), we performed a propensity score matching analysis. Genotyping and associative analyses of rs10770141, rs4680, and rs1800497 with schizophrenia were performed. Results: We analyzed 1,271 schizophrenics (male/female: 574/698; age 47.4±13.9 years) and 1,271 matched controls (male/female: 603/669; age 46.5±13.4 years). The estimated odds ratios (ORs) were 1.245 (p<0.001) for rs4680, 1.727 (p<0.0001) for rs1800497, and 1.788 (p<0.0001) for rs10770141. Double SNP analyses revealed the ORs of 2.010 (p<0.0001) for the combination of rs4680*rs1800497, 1.871 (p<0.001) for rs1800497*rs10770141, and 1.428 (p=0.068) for rs4680*rs1800497. Among the individuals with any of the three double SNP risk combinations (which accounted for 35.8% of the involved patients), the estimated OR was 2.224 (p<0.0001). Conclusions: In this validation study, the combinations of functional polymorphisms related to dopaminergic genes were associated with the development of schizophrenia. Analyzing combinations of functional polymorphisms with the control of possible confounders may provide new insights for association research.

## Introduction

Schizophrenia is a disabling chronic mental illness with the lifetime prevalence of 0.4%–1.0% worldwide [1,□2]. The pathogenesis of this disorder has been considered multifactorial, but the genetic and environmental risk factors are not fully identified [3]. Schizophrenia may also be considered heterogeneous based on dimensions including its severity, social function, long-term prognosis, sex differences, age at onset, symptomatology, and response to antipsychotic treatment [4–7]. This heterogeneity may indicate that schizophrenia could be classified into several subtypes associated with different risk genes [8–10]. It is thus important to reduce this heterogeneity by excluding confounding factors by matching the biological characteristics of individuals with schizophrenia, especially when conducting genetic association studies.

The majority of individuals with first-episode schizophrenia respond positively to antipsychotic treatment with dopamine D2 receptor (DRD2) antagonists [5]. Increased dopamine synthesis was observed in these responding patients [11]. These observations suggest that abnormal dopaminergic signaling may be involved in the pathology in a subset of schizophrenia. However, some patients with schizophrenia acquire resistance to antipsychotic treatments during their clinical course. This resistance, called dopamine supersensitivity psychosis (DSP) [12,□13], might be caused by a compensatory upregulation of DRD2 as a consequent neurobiological response to long-term antipsychotic treatment.

Based on our speculation that DSP could be thought of as a type of dopamine-related schizophrenia, we have examined the literature concerning a pathology hypothesis [14], genetic studies [15–17] and suggested treatments [13]. We recently conducted a preliminary study, and its results demonstrated that the following could indicate risks for schizophrenia [17]: the specific allele combinations of the functional single nucleotide polymorphisms (SNPs) rs10770141, rs4680, and rs1800497, which influence the activities of dopaminergic *tyrosine hydroxylase* (*TH*) gene, *catechol-O-methyltransferase* (*COMT*) gene, and *dopamine D2 receptor* (*DRD2*) gene, respectively.

In genetic association studies, control subjects have no history of mental disorders at the time point of the studies. However, the lifetime prevalence of common mental disorders was estimated as almost 30% [18], and the onsets of these disorders are distributed by age [19]. It is thus possible that some individuals who have genetic risks but have not yet reached the onset of a mental disorder are included in healthy control groups. The impact of including pre-disease subjects could also vary by their ages, suggesting a potential influence of the age distribution on the outcomes of association studies. When such studies are conducted, it is thus important to evaluate any associations between subject groups with matched clinical and/or biological features.

One approach to reduce or eliminate the effects of selection bias and confounding effects is the use of propensity score matching, which allows the design and analysis of an observational (nonrandomized) study that mimics some of the characteristics of a randomized controlled trial. Here, we conducted a secondary validation study of the large-scale age- and sex-matched case-control populations examined in our earlier research.

## Subjects and Methods

### Subjects

All of the subjects were recruited from the Honshu area of Japan (the main island of Japan), where the population falls into a single genetic cluster [20]. The diagnosis of schizophrenia was based on the Diagnosis and Statistical Manual of Mental Disorders IV (DSM-IV) criteria and confirmed by at least two experienced psychiatrists. Controls were interviewed by experienced psychiatrists to exclude any past or present psychiatric disorders. Using a subset of those subjects, we showed that the population stratification was negligible in our sample [21].

In the present study, we excluded all of the individuals recruited at Chiba University and its affiliated hospitals so that we could conduct the validation in a population that was entirely different from the subjects who participated in our previous preliminary study. All participants gave informed, written consent to join the study after receiving a full explanation of study protocols and objectives. The study was approved by the ethics committees of Chiba University, RIKEN, and all participating institutes and was conducted in accordance with the Declaration of Helsinki.

### Single nucleotide polymorphisms

We examined three SNPs, i.e., rs10770141, rs4680, and rs1800497, which showed a possible association with the onset of schizophrenia in our preliminary study [17]. Patients carrying the minor allele of rs10770141, i.e., T(+), were reported to exhibit a 30%–40% higher gene expression of *TH* compared to T(−) individuals [22]. The SNP rs4680 is a non-synonymous SNP that causes the replacement of valine with methionine at residue 158, resulting in an almost 50%–75% reduction of the enzymatic activity of COMT [23]. Carriers and non-carriers of the methionine allele are expressed as Met(+) and Met(−), respectively. For rs1800497, imaging analyses by positron emission tomography revealed that individuals with the minor allele, i.e., A1(+), had 12% less availability of striatal DRD2 than those with the major allele, A1(−) [24].

### Genotyping

SNP genotyping was performed by TaqMan SNP genotyping assays (Applied Biosystems, Foster City, CA, USA). We used a GeneAmp PCR System 9700 Dual 384-Well Sample Block Module (Applied Biosystems) for the polymerase chain reaction, and we analyzed the fluorescent signals using a 7900HT Sequence Detection System and SDS v2.4 software (Applied Biosystems).

### Statistical analyses

All data were analyzed according to the intention-to-treat principle. For the baseline variables, summary statistics were constructed by using frequencies and proportions for categorical data and means□±□standard deviations for continuous variables. We compared patient characteristics by using the chi-square test or Fisher’s exact test for categorical outcomes and t-tests or the Mann-Whitney U-test for continuous variables, as appropriate.

In order to reduce bias and a large discrepancy in the number of subjects in the groups, we performed a case-matched study by using the propensity score matching method with a ‘Greedy 5-To-1 Digit-Matching’ algorithm for patient characteristics (gender and age). Hardy-Weinberg equilibrium testing was used as a quality control for genotyping. Fisher’s exact test, crude odds ratios (OR), and 95% confidence intervals (CIs) were used to compare allele, dominant, and recessive models between groups. The significance of an association in the second screening was evaluated at the significance level of 0.05 by multiple testing using the Bonferroni correction method. All analyses were performed using SAS software, ver. 9.4 (SAS Institute, Cary, NC).

## Results

### Comparisons of the schizophrenia and control groups

We analyzed 2,012 patients with schizophrenia (1,111 men, mean□±□SD age 47.2□±□14.1 years; 901 women, age 49.2□±□14.7 years) and 2,170 matched controls (889 men, age 39.2□±□13.8 years; 1,281 women, age 44.6□±□14.1 years) from the Japanese population. We examined a total of 1,271 schizophrenia cases and 1,271 matched healthy controls.

The clinical characteristics of the participants are summarized in Table 1. As matched by propensity score, there were no significant distributional differences in sex or age between the groups. Of the 1,271 patients with schizophrenia, we included 574 males and 698 females. Their mean age at evaluation was 47.4□±□13.9 years (range 20–81 years). The mean age at disease onset was 24.9□±□13.4 years. Of the 1,271 healthy controls, we included 603 males and 669 females. Their mean age at evaluation was 46.5□±□13.9 years (range 20–86 years).

**Table 1.** Comparison of the schizophrenia and control groups

|  | **Schizophrenia**<br>**n=1271** | **Controls**<br>**n=1,271** | **p-value** |
| --- | --- | --- | --- |
| Males / females | 574 / 698 | 603 / 669 | 0.249<sup>a</sup> |
| Age at evaluation, mean (SD), yrs | 47.4 (13.9) | 46.5 (13.4) | 0.095<sup>b</sup> |
| Age range, yrs | 20–81 | 20–86 |  |
| Age at onset, mean (SD), yrs | 24.9 (13.4) |  |  |
| Onset age range, yrs | 9–73 |  |  |

There were no significant distributional differences in sex or age between the groups. The data for the age at onset were missing in 68 cases.
<sup>a</sup>By the chi-square test.
<sup>b</sup>By Student's t-test.

### Distribution of genetic risks

The distributional patterns of each genetic risk are shown in Figure 1. Within the group of schizophrenic patients, the proportions of risk carriers were 48.1% for rs4680, 61.1% for rs1800497, and 11.5% for rs10770141. The risk carriers of rs4680*rs1800497 (shown as A in Fig. 1) were 29.7%, those of rs4680*rs10770141 were 5.2% (shown as B), and those of rs1800497*rs10770141 (shown as C) were 6.9%. Note that these carriers included the patients who had all three genetic risks (shown as D, 2.9%). The patients without risk comprised 18.1%.

**Fig. 1.** For convenience, the related risk genes are listed instead of SNP IDs. The IDs for COMT, DRD2, and TH are rs4680, rs1800497, and rs10770141, respectively. The capital letters indicate the genetic combinations of rs4680*rs1800497 (A), rs4680*rs10770141 (B), rs1800497*rs10770141 (C), and rs4680*rs1800497*rs10770141 (D). Note that groups A–C also included group D.

Likewise, in the healthy control group, the risk carriers of rs4680 were 42.7%, those with rs1800497 were 47.6%, and those with rs10770141 were 6.8%. Regarding the double combination risks, group A’s percentage was 17.3%, that of group B was 3.8%, that of group C was 3.8%, and that of group D was 2.4%. The controls without risk accounted for 25.3%.

### Genetic risks

In our age- and sex-matched population, we observed significant associations of all tested rs4680, rs1800497, and rs10770141 with the onset of schizophrenia. The odds ratios were estimated as 1.245 (95%CI: 1.065–1.456, p<0.001) for rs4680, 1.727 (95%CI: 1.475–2.021, p<0.0001) for rs1800497, and 1.788 (95%CI: 1.354–2.361, p<0.0001) for rs10770141 (Fig. 2). The double SNP analyses revealed significant associations for the combinations of rs4680*rs1800497 (OR 2.010, 95%CI: 1.664–2.427, p<0.0001) and rs1800497*rs10770141 (OR 1.871, 95%CI: 1.306–2.680, p<0.001). The combination of rs4680*rs1800497 showed a tendency (p=0.068) to be associated with the onset of schizophrenia, with the OR of 1.428 (95%CI: 0.976–2.088).

**Fig. 2.** Odds ratios and 95%CI values were estimated by Fisher’s exact test. For convenience, the related risk genes are listed instead of SNP IDs. The IDs for COMT, DRD2, and TH are rs4680, rs1800497, and rs10770141, respectively.

The combination of all three SNPs showed no association with the disease onset (OR 1.240, 95%CI: 0.765–2.012, p=0.458). Within the subset of individuals with any of the three double SNP risk combinations, the disease risk was estimated as OR 2.224 (95%CI: 1.860–2.659, p<0.0001). Detailed data are provided in Supplementary Table S1.

## Discussion

We identified significant associations of the SNPs rs10770141, rs4680, and rs1800497 with schizophrenia, with ORs ranging from 1.24 to 1.79. According to our examination of the literature, the involvement of rs10770141 in the onset of schizophrenia has been suggested, but most of the prior studies failed to demonstrate a significant genetic association [25]. In addition, rs4680 has been reported to have a significant association with schizophrenia [26], but this finding was not always supported [27]. The results of studies focusing on rs1800497 also indicated that its association with schizophrenia remains controversial [28, 29].

If schizophrenia consists of multiple pathologies that are associated with different groups of genetic backgrounds, potent cofounders included in sample populations might highlight the relative importance of identifying the risk genes. In the present study we performed a propensity score matching analysis, which may increase the homogeneity between the sample groups by excluding cofounding influences of sex and age.

We observed that the risk of combinations□—□which accounted for 35.8% of the present schizophrenia patients□—□showed a significant association with schizophrenia, with the OR of 2.224. These combinations may indicate some pathological characteristics of dopaminergic transmission. The SNPs rs10770141, rs4680m, and rs1800497 are functional polymorphisms that may yield relatively high dopamine synthesis [22], rapid dopamine degradation [23], and low DRD2 density [24], respectively. Another group reported that the A1 allele of rs1800497 was associated with increased dopamine synthesis, possibly due to decreased autoreceptor function at presynaptic dopamine neurons [30].

These observations may indicate that having either or both of rs10770141 and rs1800497 could result in a rapid elevation of the synaptic dopamine concentration in response to neuronal excitation. Although COMT has not been reported to show a significant influence in the striatal cortex [31] but was shown to act predominantly in the prefrontal cortex (PFC) [32], the attenuation of dopamine signals in the PFC may lead to an increased dopamine release in the nucleus accumbens (N.Acc) (Fig. 3). Decreased activity of glutamatergic neurons at the PFC could cause less excitation of gamma-aminobutyric-acid (GABA) neurons in the ventral tegmental area (VTA), which is inhibitory against dopamine neurons projecting to the N.Acc. These functional polymorphisms might share common characteristics of increased dopaminergic neurotransmission. The synergistic trend we observed in which having multiple alleles indicated a higher risk compared to the single SNPs suggests the possibility that some patients may have abnormal dopaminergic neurotransmissions derived from these genes.

**Fig. 3.** COMT: catechol-o-methyl transferase, DA: dopamine, DLPFC: dorsolateral prefrontal cortex, DRD2: D2 dopamine receptor, GABA: gamma-aminobutyric acid, Glu: glutamate, N.Acc: nucleus accumbens, TH: tyrosine hydroxylase, VTA: ventral tegmental area.

Excessive dopamine signaling induced by a dopamine agonist [1] or an indirect agonist such as amphetamine [34] may cause an acute psychotic state. It was suggested that frequent exposure to stress could be a risk for prodromal psychotic symptoms and schizophrenia [35]. This phenomenon may be similar to the drug-induced sensitization caused by repeated exposure to dopamine agonists [36]. This sensitization is described as a progressive and long-lasting amplification of the behavioral and neurochemical response [37] and was also reported to be caused by stressors [38]. Having a combination of the risk alleles could possibly potentiate the fluctuation in synaptic dopamine concentrations, resulting in the development of a preferable environment for the sensitization.

Another significant observation in the present investigation is the even higher risk (OR 3.957, 95%CI: 2.830–5.533, p<0.001) among the subjects who were ≤40 years old, including as much as 39.2% of the disease population (data not shown). The development of prophylactic interventions especially for individuals at these younger ages with the identified risks is desired.

Indeed, the identification of individuals with genetic risks could allow us to provide them with prophylactic intervention. Assuming that there is an unstable dopaminergic state prior to the onset of schizophrenia, the recently reported association of environmental factors with vulnerability to psychosis [39] seems plausible. If so, it may be possible that even non-pharmacological treatments such as cognitive behavioral therapy could also contribute to disease prevention/treatment by strengthening an individual’s adaptability to stressors. In addition, determining the combinations of functional polymorphisms related to schizophrenia would contribute to a better understanding of the disease pathology and its further subcategorization. These advances will exert a salutary influence on clinical practice, especially for improving the accuracy of diagnoses.

This strategy may also be applicable for other multifactorial diseases including diabetes and hypertension. We believe that the possession of multiple functional genetic risks, which may indicate the same directional influences toward a pathogenic environment, would be a strong candidate as a new risk predictor. Because most large-scale association studies tend to be retrospectively conducted, statistical techniques such as the propensity score matching we used herein should be considered, in order to reduce the influence of possible cofounding factors.

Gene-environment interactions also affect the onset of schizophrenia [3,40]. In light of the current understanding of this disease, it appears that schizophrenia contains some heterogeneity in its onset mechanism. The disease state of some patients with schizophrenia may thus be more closely related to another, non-dopaminergic pathology such as glutamate- or GABA-mediated neurotransmission. Further studies are necessary to determine whether this approach can reveal associations among other factors.

