## Supplementary figures and images for "Genetic risks of schizophrenia identified in a matched case-control study"

### Fig1

**Fig. 1.** Distribution of genetic risks.

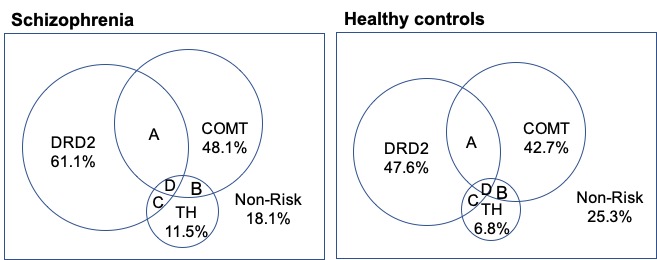

### Fig2

**Fig. 2.** Genetic risks for schizophrenia

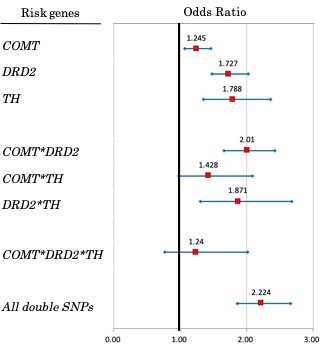

### Fig3

**Fig. 3.** Hypothetical schema of neuronal connectivity (modified from [33]).

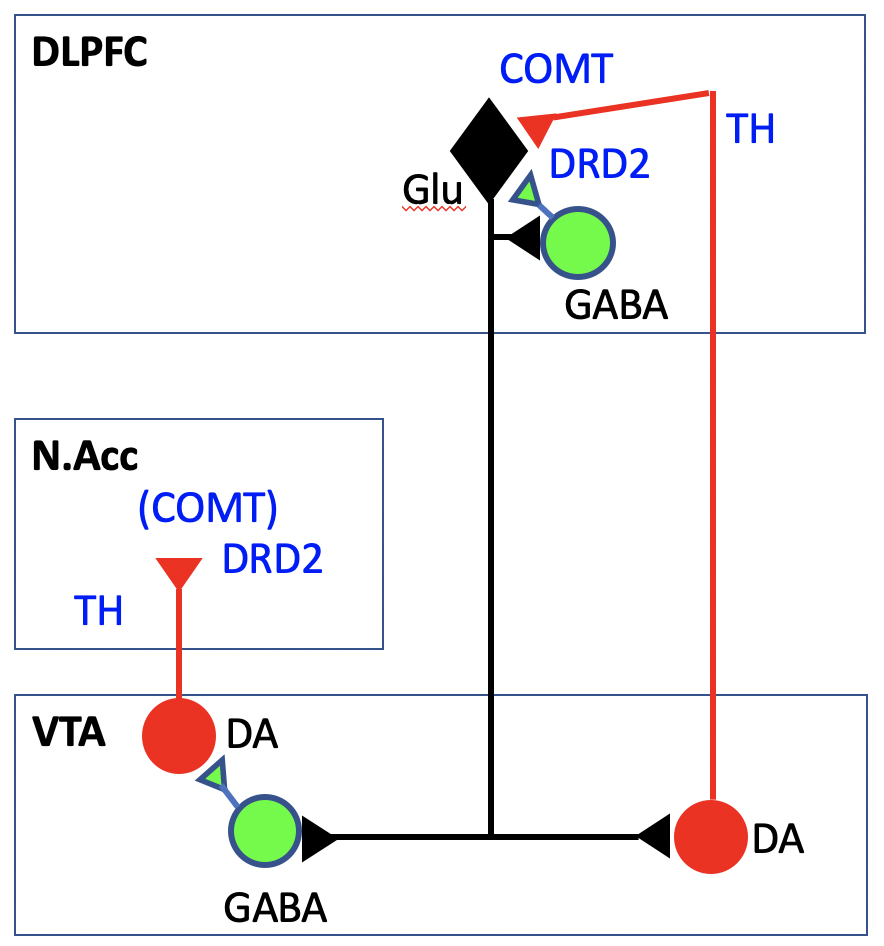
