## Supplemental Table 1 for "Genetic risks of schizophrenia identified in a matched case-control study"

**Supplementary Table S1.** Risk distributions

| Risk SNPs |  | SZ | CNT | %SZ | ％CNT | OR | lower | upper | p-value |
| --- | --- | --- | --- | --- | --- | --- | --- | --- | --- |
| COMT | Risk | 611 | 543 | 48.1% | 42.7% | 1.245 | 1.065 | 1.456 | 0.007 |
|  | Others | 658 | 728 | 51.9% | 57.3% |  |  |  |  |
| DRD2 | Risk | 776 | 605 | 61.1% | 47.6% | 1.727 | 1.475 | 2.021 | 0.000 |
|  | Others | 494 | 665 | 38.9% | 52.4% |  |  |  |  |
| TH | Risk | 146 | 86 | 11.5% | 6.8% | 1.788 | 1.354 | 2.361 | 0.000 |
|  | Others | 1125 | 1185 | 88.5% | 93.2% |  |  |  |  |
| Double |  |  |  |  |  |  |  |  |  |
| COMT*DRD2 | Risk | 376 | 220 | 29.7% | 17.3% | 2.010 | 1.664 | 2.427 | 0.000 |
|  | Others | 892 | 1049 | 70.3% | 82.7% |  |  |  |  |
| COMT*TH | Risk | 66 | 47 | 5.2% | 3.8% | 1.428 | 0.976 | 2.088 | 0.068 |
|  | Others | 1203 | 1223 | 94.8% | 96.3% |  |  |  |  |
| DRD2*TH | Risk | 87 | 48 | 6.9% | 3.8% | 1.871 | 1.306 | 2.680 | 0.001 |
|  | Others | 1183 | 1221 | 93.1% | 96.2% |  |  |  |  |
| Triple |  |  |  |  |  |  |  |  |  |
| COMT*DRD2*TH | Risk | 37 | 30 | 2.9% | 2.4% | 1.240 | 0.765 | 2.012 | 0.458 |
|  | Others | 1231 | 1238 | 97.1% | 97.6% |  |  |  |  |
| Any of three double-risk SNPs combinations | | | |  |  |  |  |  |  |
| Any Double | Risk | 455 | 255 | 35.8% | 20.0% | 2.224 | 1.860 | 2.659 | 0.000 |
|  | Others | 816 | 1017 | 64.2% | 80.0% |  |  |  |  |

The OR, 95%CI, and p-values were estimated by Fisher’s exact test. For convenience, the related risk genes are listed instead of SNP IDs. The IDs for COMT, DRD2, and TH are rs4680, rs1800497 and rs10770141, respectively. COMT, catechol-o-methyl transferase; TH, tyrosine hydroxylase; DRD2, D2 dopamine receptor; SZ, schizophrenia; CNT, control.
